# Predicting Neuroplasticity Effects of Continuous Theta Burst Stimulation with Biomarkers from the Motor Evoked Potential TMS Input-Output Curve

**DOI:** 10.1101/2025.02.20.638871

**Authors:** Shreya Parchure, Zihang Xu, Priyanka Shah-Basak, Brian Erickson, Denise Harvey, Rachel Wurzman, Darrian McAfee, Daniela Sacchetti, Olufunsho Faseyitan, Roy H. Hamilton

## Abstract

The field of neuromodulation lacks predictors of individual differences in plasticity that influence responses to repetitive transcranial magnetic stimulation (rTMS). Continuous theta burst stimulation (cTBS), a form of rTMS known for its inhibitory effects, shows variable responses between individuals, potentially due to differences in neuroplasticity. Predicting individual cTBS effects could vastly enhance its clinical and experimental utility. This study explores whether motor evoked potential (MEP) input-output (IO) parameters measured prior to neuromodulation can predict motor cortex responses to cTBS. IO curves were sampled from healthy adults by recording MEPs over a range of single pulse TMS intensities to obtain parameters including MEP_max_ and S_50_ (midpoint intensity). Subjects later received cTBS over the same location of motor cortex and their MEPs before and after stimulation were compared. Both MEP_max_ and S_50_ predicted responses, significantly correlating (p<0.05, R^2^>0.25) with individuals’ MEP changes at 10, 20, and 30 minutes after cTBS. Further, we introduced and validated an easily implementable biomarker that does not require the time-consuming sampling of full IO curve: MEP_130RMT_ (median of 10 MEPs at 130% RMT). MEP_130RMT_ was also a strong predictor of cTBS response (p<0.005, R^2^>0.3). Head-to-head comparison against a previously studied genetic biomarker of rTMS responses (BDNF polymorphism) showed that IO based predictors had a superior performance in explaining more response variability. Thus, IO curves derived prior to cTBS administration can reliably predict cTBS-induced changes in cortical excitability. This work points toward an accessible strategy for tailoring stimulation procedures in both diagnostic and therapeutic applications of rTMS, and potentially boosting response rate to other brain stimulation approaches.

**HIGHLIGHTS:**

- Baseline TMS-MEP Input-Output (IO) Curve parameters significantly predict MEP responses to M1 cTBS.
- Higher MEP_max_ at baseline predicts more robust inhibitory response to cTBS, while higher midpoint intensity (S_50_) is associated with less response.
- D We developed and validated a new biomarker MEP_130RMT_, which predicts cTBS response using just 10 baseline MEPs from single TMS pulses of 130% RMT intensity.
- Head to head comparison against BDNF genotyping shows superior performance of IO biomarkers.

## INTRODUCTION

Clinical and research applications of repetitive transcranial magnetic stimulation (rTMS) are rapidly growing due to its ability to non-invasively probe and modulate cortical activity. Since FDA approval in 2008, rTMS has become widely used as a treatment for major depressive disorder [1] obsessive-compulsive disorder [2], addiction [3], and is being explored as an intervention for other neurological and psychiatric disorders [4]. It also remains a critical research tool for elucidating the structure-function relationships in the brain related to a wide range of motor, perceptual, and cognitive abilities [5], [6], [7], and for characterizing the physiologic mechanisms that underlie cortical excitability [8] and neuroplasticity [9]. Theta burst stimulation (TBS), a modified form of rTMS, is understood to produce robust effects on cortical excitability in a fraction of the time of other rTMS protocols, making it an attractive approach for research and clinical applications. Continuous TBS (cTBS), which consists of 50 Hz bursts of stimulation pulses delivered in triplets at 5 Hz, has been associated with inhibitory aftereffects on cortical excitability [10] analogous to those from 1Hz and other inhibitory rTMS protocols. These aftereffects last from 30 minutes following 20s of cTBS (300 pulses) to 60 minutes following 40s of cTBS (600 pulses), which is longer lasting than other inhibitory TMS methods [11].

However, an emerging problem related to the use of rTMS and specifically cTBS is high inter-individual variability in physiologic responses [12], [13]. We do not yet have a clear, consistent way to determine who best responds to cTBS or who will not, which critically impedes the ability to stratify who should receive this form of intervention and limits its overall efficacy. Studies have also shown inconsistent responses to cTBS [14], which may be partially attributed to factors such as differences in age [15], time of day [10], hormone levels [16], variability in motor threshold [17], and use of substances that may alter synaptic threshold [18]. In addition, a few recent studies by our group [19], [20], [21] and others [22], [23] have also investigated neurotrophins such as brain derived neurotrophic factor (BDNF), an important factor for neural plasticity and regrowth with naturally occurring variations based on genotype, as mediators of varying responses to non-invasive brain stimulation paradigms including cTBS. These different pieces of evidence suggest that individual differences in response to cTBS may be in large part due to differences in neuroplasticity [24]. Thus, developing a quick, objective, easily translatable neuroplasticity-based predictor of responses to TMS may help to personalize individual treatments and improve group-level TMS response rates.

TMS input-output (IO) curves are one such validated measure of an individual’s cortical plasticity and neural excitability after brain stimulation [25]. IO curves measure the relationship between stimulation intensity of single TMS pulses and the amplitude of motor evoked potentials (MEPs) in the target muscle. The neurophysiological mechanism underlying IO curves is based upon the activation of cortical neurons due to single TMS pulse and propagation of signals down the corticospinal tract leading to muscle contraction in an MEP [26], and can help determine rTMS stimulation intensity [27]. Here we study individual-specific IO curves (i.e., the graph of MEP response vs. TMS intensity) that is commonly fitted using a Boltzmann sigmoidal function [28]. The graph’s slope, midpoint value *S*_50_ and plateau value MEP_max_ are independent, reliable and popular metrics of neuroplasticity [29], [30], [31]. Research shows that IO curves mirror and predict changes in cortical motor maps, such as those changes arising from due to amputation or ischemia [32], [33]. Studies of IO curves in conjunction with motor cortex rTMS show these IO parameters are a good index of neural excitability, and they are commonly used to demonstrate cTBS-induced modulation of cortical excitability or LTD-like neuroplastic responses [13]. Specifically, IO curves with steeper slopes are associated with inhibitory 1Hz rTMS effects, while decreases in the *S*_50_ and MEP_max_ values are associated with the excitatory effects of 20Hz rTMS [30].

In pursuing this experiment, we aimed to develop a biomarker that is based upon individual differences in cortical physiology, to predict differences in response to cTBS. The ability to predict individual responses to cTBS and other neuromodulation approaches could, in turn, allow for further optimization of therapeutic and experimental TMS interventions. Here, we hypothesized that IO curves, sampled by recording MEPs over a range of sub- and supra-threshold single pulse TMS intensities, can be used to predict MEP changes induced by cTBS subsequently delivered to the same region. Since IO curves are indicative of neuroplastic responses to TMS, we predicted that individuals exhibiting a baseline IO curve with parameters indicative of greater excitability (i.e., steeper slope, lower S_50_ midpoint, higher MEP_max_ plateau) would show expected inhibitory cTBS effects on MEPs. Given the high variability in responses to cTBS [12], [14], we expected that some subjects would show minimal MEP changes or even paradoxical excitatory MEP changes. We also hypothesized that these individuals would have IO curve parameters suggestive of less cortical excitability. We also tested the performance of our measure relative to other research biomarkers of neural plasticity and excitability that have been associated with TMS responses - such as polymorphisms in the BDNF gene [22]. Leveraging recent studies showing redundancy between certain IO curve parameters with the median MEP at a single supra-RMT stimulation intensity [31], we then sought to develop and benchmark a simple biomarker that could be quickly assessed without sampling full IO curve for increased translational potential. This work may provide cortical excitability-informed biomarkers predictive of neural responses to cTBS. More broadly, the ability to successfully predict the effects of TMS based on physiological markers would help to optimize and personalize TMS treatments.

## METHODOLOGY

### Experimental Overview

The experiment consisted of two sessions (Refer to Figure 1). The initial visit consisted of recording the input-output (IO) curves after determining the TMS location within the left motor cortex used to elicit MEPs, and the TMS intensity based upon motor thresholds. The second visit, occurring 1-7 days later, consisted of experimental manipulation of the same cortical site using cTBS for 40 sec (600 pulses). The changes of cortical excitability in response to cTBS were indexed by collecting 30 MEPs each at baseline as well as 0, 10, 20, and 30 minutes after cTBS.

**Figure 1.**
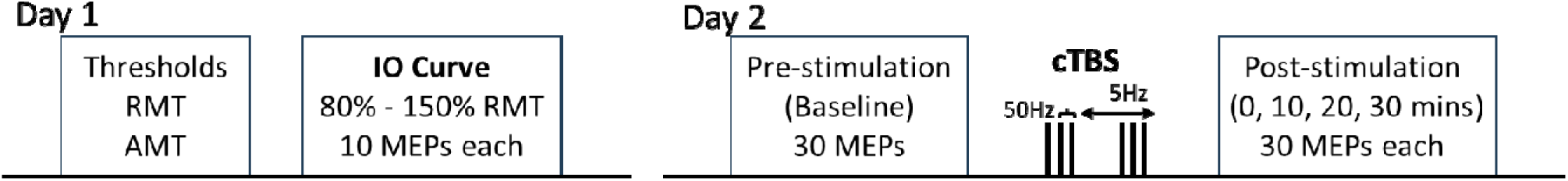
Schematic of experimental design. Input-output (IO) curve was collected as the graph of 10 MEPs each from 10 TMS intensities at 80-150% of individual RMT. At a later day, 30 MEPs were collected both before and at 10 min intervals after cTBS to the same stimulation site. RMT = Resting motor threshold, AMT = active motor threshold, MEP = motor evoked potential.

### Participants

Subjects included 21 healthy right-handed native English speakers, ages 18-35 (10 female and 11 male). Participants had no contraindications to TMS [34] and no neurological or psychiatric disorders. All experimental protocols were conducted in accordance with IRB guidelines at University of Pennsylvania.

### TMS Site Determination and Resting Motor Threshold (RMT)

Participants’ T1-weighted MRI scans were uploaded to the Brainsight® Neuronavigation system (Rogue Research, Montreal) and were used to identify and store the optimal TMS coil position over the left primary motor cortex for eliciting motor evoked potentials (MEPs) from the right first dorsal interosseous (FDI) muscle. Single pulses of TMS with a monophasic waveform were administered to the primary motor cortex of the left hemisphere using a Magstim 200^2^ Stimulator with a 70mm figure-eight coil (Magstim Co., Whitland, Dyfield, UK). To assess MEP amplitudes evoked by TMS, electromyographic (EMG) activity was recorded using surface electrodes spanning the belly of the first dorsal interosseous (FDI) muscle of each patient’s right hand, with ground electrode placed along the wrist. In line with accepted methods [10], [11], resting motor threshold (RMT) was defined as the minimum TMS pulse intensity required to elicit MEPs with peak-to-peak amplitudes of at least 50 μV in 5 of ten consecutive trials, while participants rested their hands on their lap or on a pillow.

### Input-Output (IO) Curve Measurement

IO curves were sampled for each subject by measuring 10 MEP responses to single TMS pulses delivered over 10 stimulation intensities from 80% to 150% of individual RMT, or to maximum stimulator output if smaller than 150% RMT. MEPs were collected using the standard EMG techniques described above. RMT and IO curves were collected on the same day, while cTBS and MEP changes from it were collected in a second session on a different day.

### Continuous Theta Burst Stimulation (cTBS) Protocol

Continuous theta burst stimulation (cTBS) was administered with a biphasic waveform using a Magstim SuperRapid^2^ Stimulator (Magstim Co., Whitland, Dyfield, UK). Subjects received cTBS over the hand representation of the left primary motor cortex; this was the same site used to determine RMT, AMT and IO curves. CTBS consisted of a continuous delivery of 50 Hz triplets of TMS pulses at 5 Hz for a total of 600 pulses (∼40s). Intensity of cTBS was set to 80% of active motor threshold (AMT), defined as the minimum pulse intensity required to produce MEPs with peak-to-peak amplitudes of at least 200 μV in 5 of ten consecutive pulses while participants contracted the FDI muscle at 20% of the maximum voluntary contraction. To ensure 20% contraction of the FDI, participants were first asked to contract with maximal force while motor activity was recorded with EMG. Subjects then practiced pushing with sufficient force to elicit 20% of the maximal EMG signal. The same biphasic stimulator was used to determine aMT and administer cTBS.

### Pre and Post cTBS Motor Evoked Potentials (MEPs)

In order to index cortical excitability before vs after cTBS, 30 MEP samples from single-pulse TMS pulses were obtained at baseline before cTBS, and post-stimulation at 0, 10, 20 and 30 minutes. The TMS pulses used to elicit MEPs were delivered at 120% RMT with an inter-stimulus interval of 6s and 6% random jitter. Neuro-navigation guided targeting was used to ensure the pre- and post-cTBS MEPs were obtained from the same cortical site and with the same TMS coil position as used during the MEP acquisition for RMT, AMT, and IO curves on prior days. Participants were seated in a chair, with their arm and hand from which MEPs were elicited resting on their lap or a pillow. MEP peak-to-peak amplitudes were recorded with the same electromyography (EMG) methods as used in RMT and IO curve sampling. Signals were amplified and band-pass-filtered between 20 and 2000 Hz, digitized (sample-rate 5 kHz), and stored for off-line analysis using SIGNAL software (Cambridge Electronic Devices, Cambridge, UK).

### BDNF Genotype Analysis

Genomic DNA from human saliva samples was collected in Oragene® DNA collection kits and then isolated using the prepIT.L2Preagent (cat # PT-L2P-5, DNA Genotek Inc, Canada) and precipitated with ethanol according to manufacturer’s instructions. The DNA samples were genotyped for BDNF (the single nucleotide polymorphism rs6265) using the TaqMan SNP Genotyping Assay (C_11592758_10) designed by Thermo Fisher Scientific. Primers and probes were mixed with TaqMan® Universal PCR Master Mix (Thermo Fisher Scientific), and 4.5 μL genomic DNA (2.5 ng/ μL) was transferred in duplicate to a plate with each well containing 5.5 μL of the PCR mixture. PCR reaction was performed following standard protocols followed by allele discrimination by post-PCR plate reading and data processing using the ViiA™ 7 Software (Thermo Fisher Scientific).

### Input-Output Curve Fit and Parameter Calculation

Each stimulation intensity at which 10 MEPs were obtained, was expressed as a percentage of the subject’s RMT. Each person’s stimulus-response (IO) curve was then calculated based on the Boltzmann sigmoidal fit relationship between stimulation intensity (% RMT, independent variable) and MEP amplitude (mV, dependent variable) [35], [36]. Refer to Figure 2 for a schematic of IO curve parameters. The sigmoid curve equation is as follows:

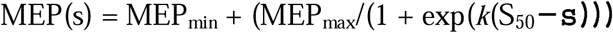

**Figure 2.**
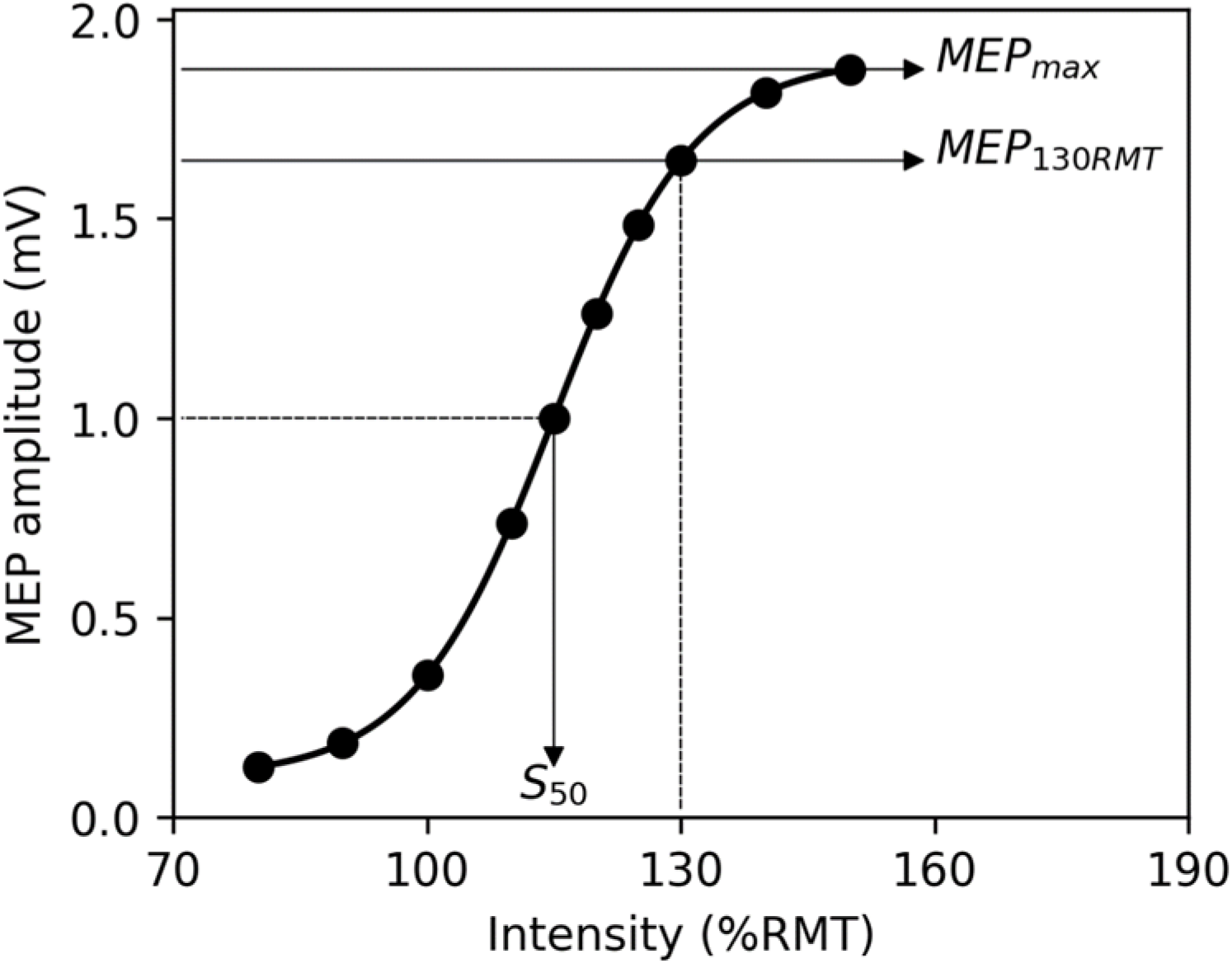
Input-Output (IO) Curve Schematic. Parameters of the IO curve studied were: S_50_= TMS intensity at the midpoint MEP value, MEP_max_= Plateau maximum MEP amplitude, MEP_130RMT_= Median MEP amplitude at 130% RMT (new biomarker created for and studied in this research).

where MEP(s) is the MEP amplitude produced by single TMS pulse of stimulation intensity s, MEP_max_ is the maximum MEP amplitude plateau and MEP_min_ is the minimum MEP amplitude, S_50_ is the stimulus intensity needed to obtain 50% of MEP_max_ amplitude, and *k* is the growth rate/ slope parameter of the sigmoid function, i.e., the global slope of the function.

Here, we focused on MEP_max_ and S_50_ which are commonly used IO properties well characterized to be independent of each other [28], [37].

The curve fitting functionality in Python was used to obtain the IO curve equation and its parameters. Sigmoidal fit was calculated using commonly employed stochastic methods with 30,000 iterations upon a standard seed guess for parameters (S_50_: 120% RMT +/- 20%, MEP_min_: 0 +/- 0.1, MEP_max_: median MEP from 150% RMT with SD, k: 1 +/- 1) and manual check to ensure stability and real world constraints. Further, a manual inspection of residuals was conducted to ensure the goodness of fit for all subjects’ IO curves. Each IO parameter and standard deviation per subject was computed in this manner.

### Computation of new Biomarker MEP_130RMT_

Further, we introduced and tested a new biomarker (MEP_130RMT_), defined as the median MEP amplitude from 10 trials at a single stimulation intensity from the IO curve – here, at 130% RMT. Such a metric does not require full IO curve sampling to determine its value, yet it may capture characteristics of the corticospinal excitability from IO curve parameters [31] and has been shown to be stable across time [38]. This new candidate biomarker, MEP_130RMT_ would be most directly correlated with MEP_max_ and inversely correlated with S_50_ based on mathematical derivation from the Boltzmann formula [39]:

Assuming MEP_min_ approaches 0 and using known correlations MEP_130RMT_ ∝ MEP_max_ and MEP_130RMT_ ∝ *k* [31] then,

MEP_130RMT_∝ constant x MEP_130RMT_ / (1 + exp(constant x MEP_130RMT_(S_50_ – 130)) Which is solvable for MEP_130RMT_ ∝ 1/S_50_

Based on above formulation, we expect to observe inverse relationship between MEP_130RMT_ and MEP changes due to cTBS as between S_50_ and MEP changes

### Statistical Analysis of MEP Changes after cTBS

We first normalized the MEP distribution by taking their natural logarithm (lnMEP). We then calculated the ‘Delta’ or change at each post-cTBS time point (P0, P10, P20, P30) from median baseline lnMEP. Positive Delta lnMEP values indicate an excitatory response while negative Delta lnMEP values indicate the expected inhibitory response to cTBS stimulation. For each post-cTBS time point, subjects were scored as responders (i.e. showing significant inhibition of MEPs compared to baseline), non-responders (i.e. showing no significant MEP changes from baseline) or paradoxical responders (i.e. showing significant MEP excitation response compared to baseline) based on a 1-way ANOVA analysis. Statistical analyses and figure generation were performed using R, Python and Excel.

### Testing for Predictors of cTBS Responses

We used ordinary least squares regression across subjects to test whether an individual’s candidate biomarker - S_50_, MEP_max_, MEP_130RMT_, and BDNF polymorphism status – would be predictive of their lnMEP changes at each time point after cTBS stimulation. The p-values for each comparison were adjusted using FDR correction for multiple comparisons.

To evaluate how a binarized version of the covariates predicts lnMEP responses in interaction with time point, we also used linear mixed effects regression models implemented in the lme4 package of R version 4.3.1 [40]. Models contained trial-level lnMEP data as the expected outcome, with input variables as each biomarker, time point, TMS stimulation parameters, and by-trial random intercept. Such a linear mixed-effects modeling approach is well suited for hierarchical biological data with more than one source of variability, to assess the relationship between a candidate independent variable and the expected outcome while adjusting for relationships of other variables with the outcome. Here, we used linear mixed effects including random effects such as the by-trial intercept which accounts for differences due to repeated MEPs. We also assessed the effect of each biomarker as a single variable and the interaction between two variables. Analyses were conducted on 3664 MEPs, which were natural log-transformed to ensure a normal distribution (lnMEP).

## RESULTS

### 1. High Variability of MEP Responses to cTBS

At the group level, there was no significant decrease in lnMEPs after cTBS, as seen through a one-way ANOVA assessing group-level changes from baseline in lnMEPs which showed no significant differences across time point (df=4, SS=0.791, MSE= 0.198, F=0.1997, p=0.938). Refer to Figure 3. A high variability in responses was present at the group level, where Mean ± SD lnMEP change from baseline at 0 minutes (immediate post cTBS) was -0.25 ± 0.7; at 10 minutes was -0.17 ± 0.6; 20 minutes was -0.22 ± 0.6; and at 30 minutes was -0.14 ± 0.7.

**Figure 3.**
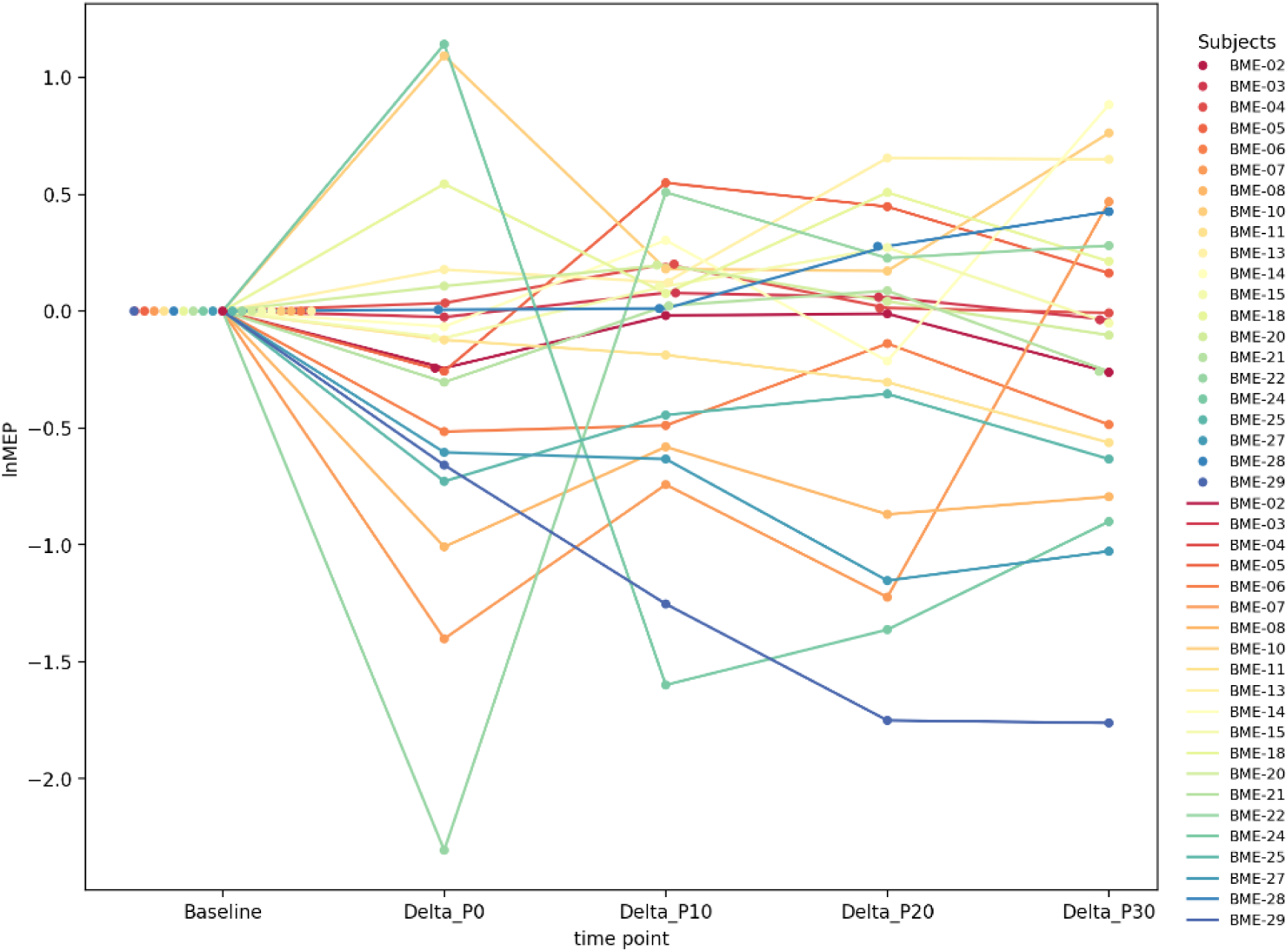
No significant group-level changes after cTBS, however individual subjects do show significant changes in excitatory and inhibitory directions as well as no significant changes. Note that all lnMEPs are normalized to individual baseline value for clarity across varying scales.

However on a within-individual basis, we observed significant changes in MEP responses after cTBS based on one-way ANOVA. A high degree of variability existed, with 8 people showing an inhibitory change post-cTBS, 7 having an excitatory MEP increase, and 6 with no significant response across 0-30 minutes after cTBS. Refer to Supplementary Table 1.

### 2. IO Curve Parameters S_50_ & MEP_max_ Predict MEP Responses to cTBS

Two independent parameters of the IO curve, midpoint intensity (S_50_) and peak plateau of th sigmoid (MEP_max_), when binarized using a median split, significantly predicted MEP response to cTBS, based on comparison of linear mixed effects models without and with interaction of these IO parameters with Time (Refer to Table 2).

**Table 1:** Individual subjects’ demographic and stimulation parameters.

| Subject ID | Sex | Race | RMT | AMT | BDNF SNP |
| --- | --- | --- | --- | --- | --- |
| BME02 | Male | AA | 53 | 45 | Val |
| BME03 | Female | C | 48 | 35 | Val |
| BME04 | Male | C | 59 | 59 | Val |
| BME05 | Female | C | 44 | 49 | Met |
| BME06 | Male | Other | 55 | 56 | Val |
| BME07 | Male | C | 50 | 44 | Val |
| BME08 | Male | C | 41 | 45 | Met |
| BME10 | Female | C | 44 | 46 | Val |
| BME11 | Female | C | 52 | 55 | Met |
| BME13 | Male | C | 74 | 74 | Val |
| BME14 | Female | A | 50 | 50 | Met |
| BME15 | Male | A | 53 | 60 | Met |
| BME18 | Female | C | 60 | 65 | Val |
| BME20 | Male | H | 58 | 62 | Met |
| BME21 | Female | C | 56 | 80 | Val |
| BME22 | Female | AA | 50 | 50 | Val |
| BME24 | Male | AA | 62 | 72 | Val |
| BME25 | Male | A | 53 | 69 | Met |
| BME27 | Female | AA | 71 | 62 | Val |
| BME28 | Male | A | 48 | 54 | Val |
| BME29 | Female | H | 46 | 53 | Val |

Key: RMT = resting motor threshold, AMT = active motor threshold, BDNF = brain derived neurotrophic factor, SNP = single nucleotide polymorphism, Val = homozygous Val/Val (n=14), Met = presence of Methionine as either Met/Met homozygote (n=3) or Val/Met heterozygote (n=4)

**Table 2:** Linear Mixed Effects Models to Compare Biomarkers.

| Model | logLik | Dev | $\chi^2$ | df | p-value |
| --- | --- | --- | --- | --- | --- |
| lnMEP ~ Time + (1 \| Subj_ID) + (1 \| Trial) | -4678.0 | 9356 |  |  |  |
| lnMEP ~ **S<sub>50</sub>** *Time + (1 Subj_ID) + (1 Trial) | -4612.5 | 9225 | 131 | 5 | < 0.001 |
| lnMEP ~ **MEP<sub>max</sub>** *Time + (1 Subj_ID) + (1 Trial) | -4659.2 | 9319 | 37.5 | 5 | < 0.001 |
| lnMEP ~ **MEP<sub>130RMT</sub>** *Time + (1 Subj_ID) + (1 Trial) | -4628.4 | 9257 | 99.2 | 5 | < 0.001 |
| lnMEP ~ **BDNF** *Time + (1 Subj_ID) + (1 Trial) | -4672.5 | 9345 | 11 | 5 | 0.052 |

The first row represents the base model, which includes covariates only. Subsequent rows show model comparison results after adding the fixed effect of interest highlighted in bold. All models were compared to the base model. Key: logLik, log-likelihood; $\chi^2$ , chi-squared test statistic; Dev, deviance; df, degrees of freedom; lnMEP, log-transformed MEP amplitude; Time, time point (Baseline vs. 0, 10, 20, or 30 min post-cTBS); BDNF, brain-derived neurotrophic factor (Val66Val vs. Val66Met); (1 | Subj\_ID), random effects structure representing a by-participant random intercept; (1 | Trial), random effects structure representing a by-trial number random intercept.

**Table 2:**
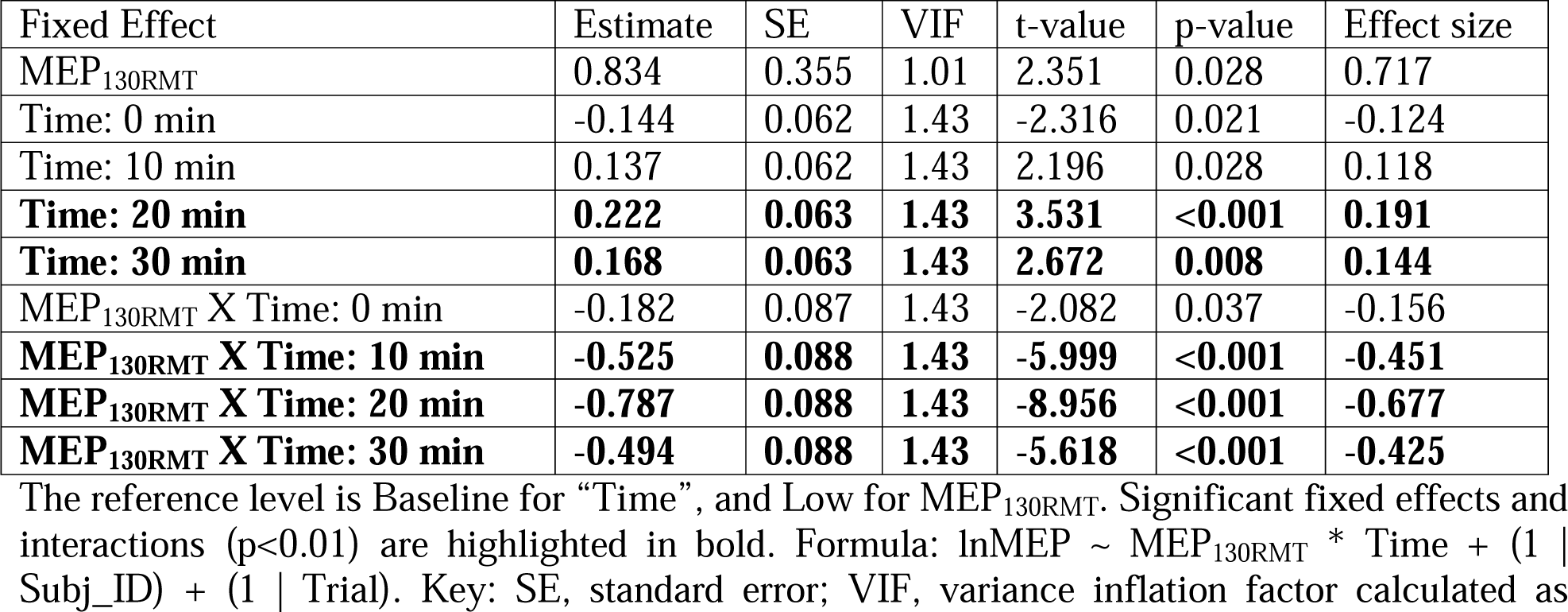
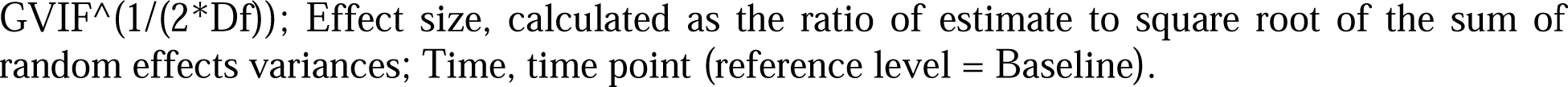
Linear Mixed Effects Model Coefficients and Associated Test Statistics.

Ordinary least squares regression between each person’s median lnMEP change from baseline vs their S_50_, showed a significant positive correlation at 10 minutes (coefficient=0.0369, SE=0.012, t=2.965, p=0.008, R^2^=0.316), 20 minutes (coefficient=0.0517, SE=0.014, t=3.726, p=0.001, R^2^=0.422), but is not significant at 0 minutes (coefficient=-0.0031, SE=0.021, t=-0.150, p=0.882, R^2^=0.001) nor 30 minutes (coefficient=0.0301, SE=0.017, t=1.819, p=0.085, R^2^=0.148) after cTBS. Refer to Figure 4.

**Figure 4:**
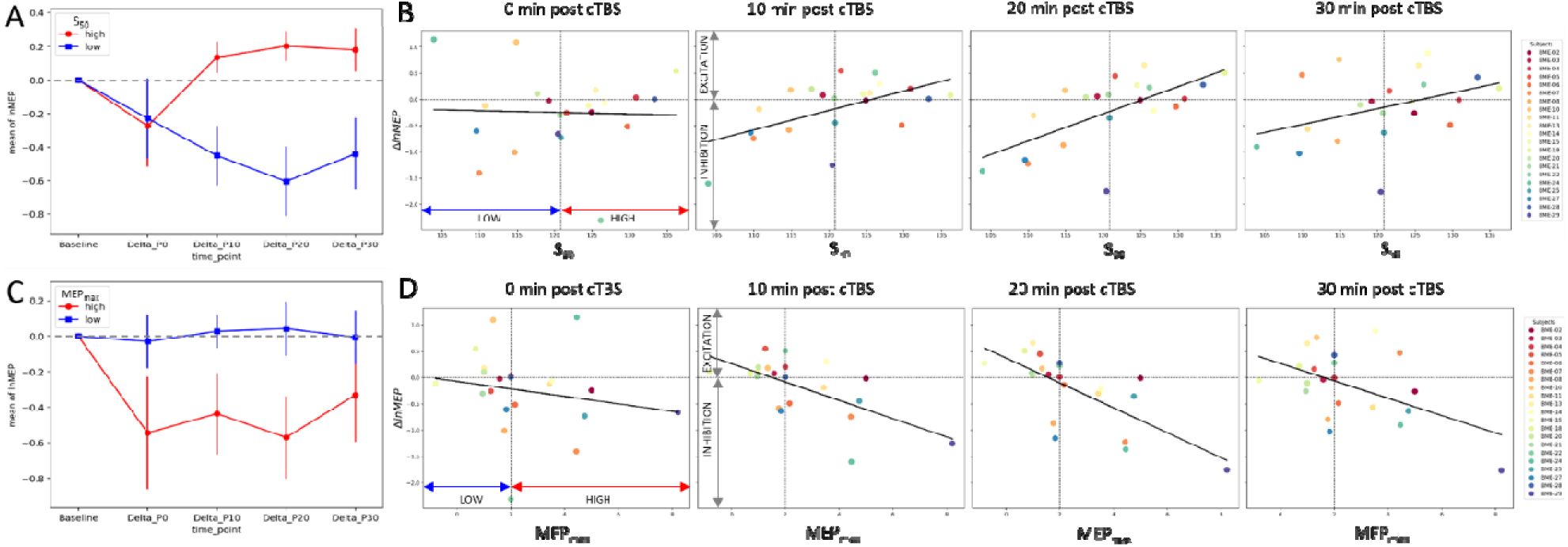
IO Curve Parameters Predict cTBS Responses (A) Difference in median LnMEPs from baseline at 0, 10, 20, and 30 minutes post-cTBS for patients with low S_50_ (blue) vs high S_50_ (red). Error bars reflect Standard Error. (B) Regression between S_50_ and change in LnMEPs from baseline at 0, 10, 20, and 30 minutes post-cTBS. (C) Difference in median LnMEPs from baseline at 0, 10, 20, and 30 minutes post-cTBS for patients with low MEP_max_ (blue) vs high MEP_max_ (red). Error bars reflect Standard Error. (D) Regression between MEP_max_ and change in LnMEPs from baseline at 0, 10, 20, and 30 minutes post-cTBS.

Regression between each subject’s MEP_max_ and their delta lnMEP changes shows significant negative correlation at post-cTBS times 10 minutes (coefficient= -0.1737, SE= 0.049, t= -3.546, p= 0.002, R^2^=0.398), 20 minutes (coefficient= -0.2364, SE= 0.054, t= -4.389, p< 0.001, R^2^=0.503), and 30 minutes (coefficient= -0.1647, SE=0.065, t= -2.537, p= 0.020, R^2^=0.253) but not significant at 0 minutes(coefficient= -0.0708, SE=0.086, t= -0.824, p= 0.420, R^2^= 0.034). Refer to Figure 4.

### 3. Median MEP from single TMS intensity as an Accessible Biomarker representing IO curves

We created and tested a novel biomarker based on median MEP from 130% RMT (MEP_130RMT_), and found this is also a significant predictor of MEP changes from baseline at following times after cTBS: 10 minutes (coefficient= -0.224, SE= 0.037, t= -6.068, p<0.001, R^2^=0.660), 20 minutes (coeff=-0.2749, SE=0.042, t=-6.474, p<0.001, R^2^=0.688), and 30 minutes (coeff=-0.2008, SE=0.059, t=-3.413, p=0.003, R^2^=0.380), but is not significant at 0 minutes (coefficient = -0.0331, SE = 0.087, t= -0.382, p=0.707, R^2^= 0.008). Refer to Figure 5. MEP_130RMT_, when binarized based on median split from clustering, is also a significant predictor of responses in interaction with Time point, as seen through linear mixed effects modeling. Refer to Table 2.

**Figure 5:**
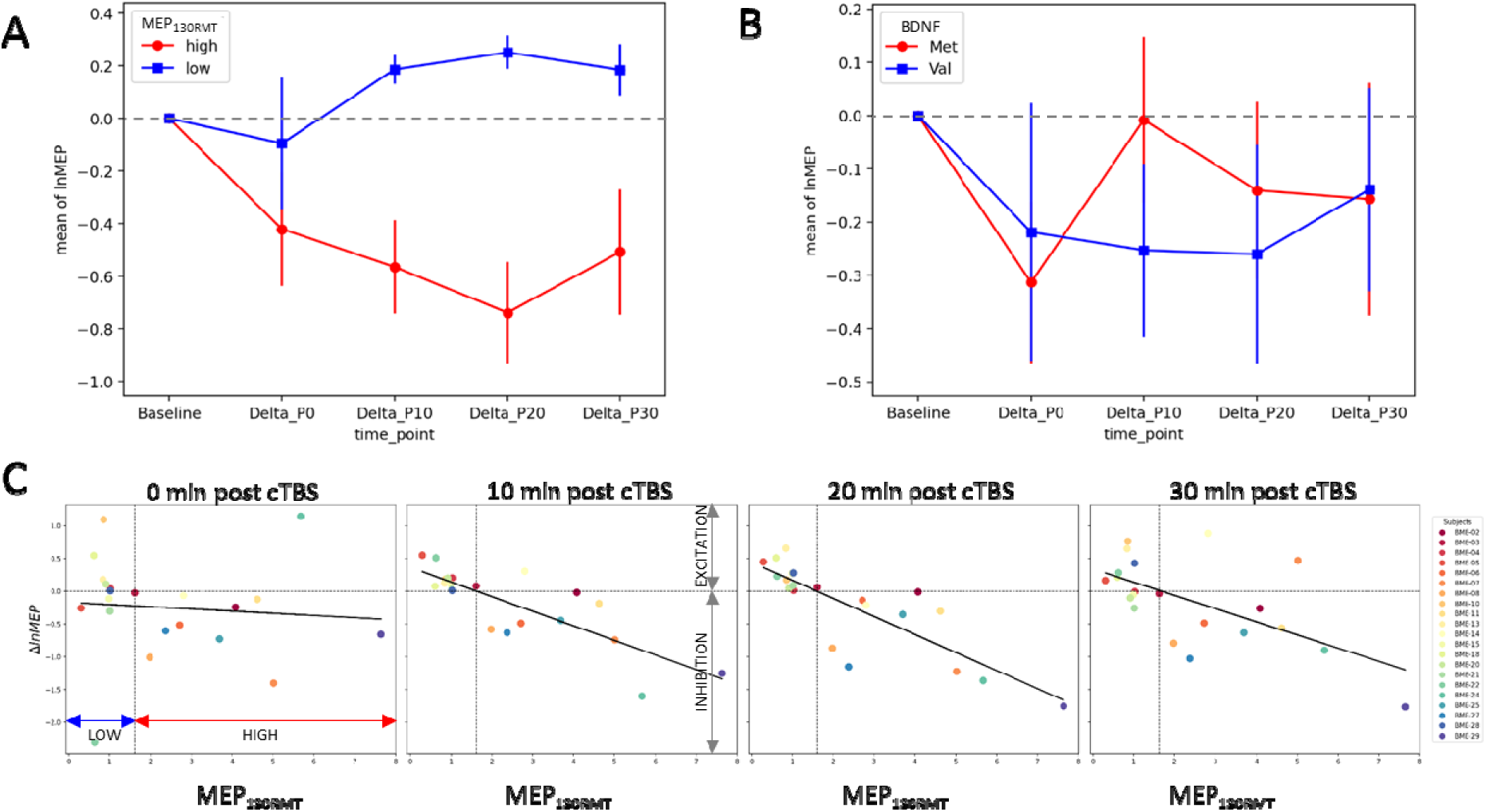
Comparison of Novel IO Biomarker (MEP_130RMT_) vs Genetic Biomarker (BDNF) in Stratifying cTBS Responders. (A) Difference in median LnMEPs from baseline at 0, 10, 20, and 30 minutes post-cTBS for patients with low MEP_130RMT_ (blue) vs high MEP_130RMT_ (red). Error bars reflect Standard Error. (B) Difference in median LnMEPs from baseline at 0, 10, 20, and 30 minutes post-cTBS for patients with BDNF Val66Val genotype (blue) vs high BDNF Val66Met polymorphism (red). Error bars reflect Standard Error. (C) Regression between MEP_130RMT_ and change in LnMEPs from baseline at 0, 10, 20, and 30 minutes post-cTBS.

### 4. IO based predictors outperform stratification using BDNF genotype in predicting TMS effects

Using linear mixed effects models to test whether an individual’s BDNF genotype (binarized by either presence of the Val66Met polymorphism or not) predicts lnMEP changes from baseline after cTBS, showed no significant effects of the BDNF interaction with Time point. Refer to Table 3 and Figure 5. Comparing BDNF with the other IO based predictors by incorporating all within one linear mixed effects model without time interaction of each showed that the binarized S50, MEPmax, and MEP130RMT accounted for significantly more lnMEP variability than BDNF. Further, the fixed effects of predictors alone at baseline or at all times combined are not significant, rather it is the interaction with time (i.e., specifically the responses to cTBS) which is significantly explained by the IO based parameters. Refer to Table 4.

**Table 3:** Linear Mixed Effects Model Coefficients and Associated Test Statistics.

| Fixed Effects | Estimate | SE | VIF | t-value | p-value | Effect size |
| --- | --- | --- | --- | --- | --- | --- |
| BDNF | 0.171 | 0.388 | 1.012 | 0.441 | 0.663 | 0.144 |
| **Time: 0 min** | **-0.231** | **0.077** | **1.740** | **-2.99** | **0.003** | **-0.194** |
| Time: 10 min | -0.231 | 0.077 | 1.740 | 0.480 | 0.631 | -0.194 |
| Time: 20 min | -0.093 | 0.078 | 1.740 | -1.188 | 0.235 | -0.078 |
| Time: 30 min | -0.077 | 0.078 | 1.740 | -0.987 | 0.324 | -0.065 |
| BDNF X Time: 0 min | -0.005 | 0.094 | 1.742 | -0.055 | 0.956 | -0.004 |
| **BDNF X Time: 10 min** | **-0.250** | **0.094** | **1.742** | **-2.648** | **0.008** | **-0.21** |
| BDNF X Time: 20 min | -0.130 | 0.095 | 1.742 | -1.374 | 0.170 | -0.109 |
| BDNF X Time: 30 min | -0.008 | 0.095 | 1.742 | -0.083 | 0.934 | -0.007 |

Reference levels are Baseline for “Time” and Val66Val for “BDNF.” Significant fixed effects and interactions ( $p < 0.01$ ) are highlighted in bold. Formula: $\ln\text{MEP} \sim \text{BDNF} * \text{Time} + (1 | \text{Subj\_ID}) + (1 | \text{Trial})$ . Key: BDNF, brain-derived neurotrophic factor (reference level = Val66Val); SE, standard error; s2, random effect variance; VIF, variance inflation factor calculated as $\text{GVIF}^{1/(2 * \text{Df})}$ ; Effect size, calculated as the ratio of estimate to square root of the sum of random effects variances; Time, time point (reference level = Baseline).

**Table 4:**
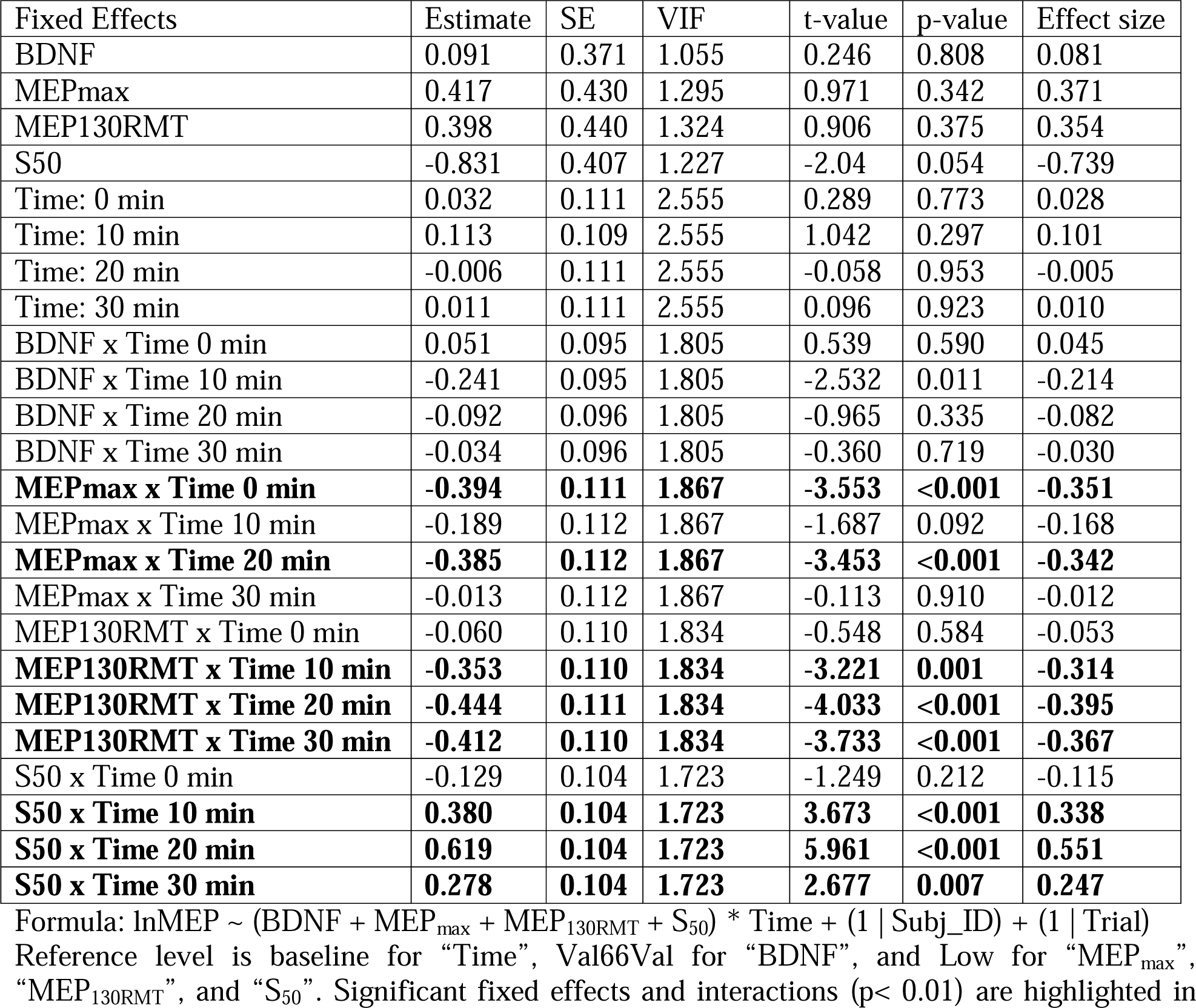
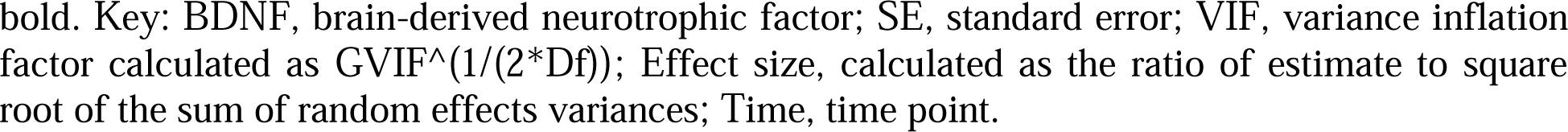
Linear Mixed Effects Model Comparing all Predictors: Coefficients and Test Statistics.

## DISCUSSION

In this study, we showed that TMS-MEP input-output (IO) curve metrics are significant predictors of the MEP change motor responses to cTBS. We were able to stratify subjects into groups of responders showing MEP inhibition vs non-responders to cTBS, using the binarized IO parameters of MEP_max_ and S50 midpoint intensity. We also created and validated a faster biomarker for translation, MEP_130RMT_, which was also significantly correlated with MEP change and was successfully used to stratify subjects. Finally, compared to more established biomarkers such as BDNF polymorphism, these IO curve based predictors are more significantly predictive of cTBS-induced changes. We found that IO curves derived prior to the administration of cTBS can serve as a reliable predictor of post-stimulation MEP responses after cTBS. Further we introduce a reliable biomarker MEP_130RMT_ that integrates into existing TMS protocols and provides a robust stratification of responders without the need for time consuming sampling of full IO curves. This addresses a major problem in the field of inter-individual high variability in responses to rTMS [14], [41], [42], while also opening the door to mechanistic understanding of cortical excitability as it relates to TMS responsiveness.

We use IO curves here as an extensively validated metric of corticospinal excitability [30], [35]. IO curves have previously been used as biomarkers of other types of neurological responsiveness such as in cortical mapping [33], [43]. It is also generally accepted that cTBS to the motor cortex produces a LTD-like neuromodulation effect by mediating the excitability of this corticospinal tract through NMDA receptors and other mechanisms [25], [44], [45]. Therefore, we expected that the IO curve index of an individual’s corticospinal tract excitability would be highly related to their responsiveness to modulation of this excitability. Most prior studies have focused on slope, a basic characteristic of the IO curve, as it relates to corticospinal excitability [46], [47] and individual motor ability such as pinch force strength [37]. However, we used more nuanced IO parameters such as S_50_ and MEP_max_ which still have known relations to IO slope [29], [31].

The slope of the IO curve has been used in relation with brain stimulation in a few studies, which suggest that steeper IO slopes correlate positively with MEP facilitation due to iTBS [43]. Extensive mapping and validation of IO parameters further shows that people with a steeper slope also have a lower S_50_ midpoint intensity [31], [35]. Thus an IO curve with a steeper slope (or lower S_50_ threshold) would be indicative of a rapidly responsive corticospinal tract and may explain those individuals’ greater responsiveness to TMS. Here, our result of the lower S_50_ group being highly responsive with strong MEP inhibition due to cTBS fits in with the literature. Further extending to other forms of brain stimulation, steeper IO recruitment curves and a higher curve ceiling (MEP_max_) have been somewhat associated with increased pinch force after tDCS stimulation over the motor cortex due to increased excitability to neuromodulation [37]. This also fits with our results of the higher MEP_max_ group showing the strong expected MEP inhibition in response to cTBS. Overall, our results also extend prior findings to show that these various IO parameters may be predictors of responsiveness in general across multiple types of TBS, rather than associated with just motor excitation or inhibition specifically.

While RMT is commonly collected prior to stimulation for purposes of determining rTMS intensity [27], sampling the full IO curve is time consuming and is not typically pursued in clinical contexts [48]. We reasoned that a less cumbersome biomarker would be more useful clinically. Our parameter MEP_130RMT_ was specifically proposed and validated with these concerns in mind, because it can be quickly sampled in only 10 MEPs after RMT determination (which is standard before all research and clinical TMS protocols). Our choice of this new biomarker MEP_130RMT_ was also backed up by prior study of redundancy and relation between IO parameters, wherein median MEP at 140% RMT is related to MEP_max_ [31]. However, we chose 130% instead because some subjects had such a high RMT that 140% would exceed the maximum stimulator output. Prior study suggests this is a highly reliable biomarker across time similar to IO curve reliability [38]. This circumvents some major barriers to use of full IO curves being time consuming or limiting the maximum doses of TMS pulses allowed in a day. Our study has validated that MEP_130RMT_ has a comparable predictive value as S_50_ or MEP_max_ yet it is also faster than those which would require the full IO curve to be sampled.

Some recent work has shown that neuroplasticity as assessed by polymorphisms in the gene coding for BDNF, associated with neural regrowth and plasticity [22], [49], impacts responses to brain stimulation including cTBS-related MEP suppression responses specifically [20], [21], [23]. We therefore sought to employ BDNF as a biomarker associated with variability in these same motor cortex cTBS protocols and compare its predictive value to that of our IO based physiologic markers. Here we went a step beyond typical predictor studies by head-to-head compared our new IO curve biomarkers directly against a previously uncovered biomarker of cTBS responses: BDNF polymorphism from genetic testing. Our results showed that IO based parameters are significantly better predictors than BDNF. We expect that BDNF is a less significant predictor amongst our small sample size for a genetic test, which may be further attributable to less Val/Met polymorphism carriers in this healthy volunteer group to be sufficiently powered for significance. However, the BDNF results are in line with prior studies [19], [22] in that the directional effects of MEP suppression or non-response are present for the same groups and even have the same variation over time. This suggests a related but more direct mechanism of IO as relates to rTMS effects. Future studies should explore the combination of BDNF and IO incorporating two distinct sources of neuroplasticity and neuronal excitability together as potential predictors of response to neuromodulation. Our work also underscores the relative advantages of these methods over each other, in that genetic testing takes more time and is costlier than an approximately 1-minute assessment of our MEP_130RMT_ biomarker, which can be obtained immediately and not just several days before TMS delivery.

We intentionally recorded a high degree of variation in cTBS responses, which is not uncommon in this field [14], [24]. In this study, having this high variability across MEP samples benefits the research question of identifying biomarkers, and shows that the stratification provided by binarized IO parameters is valid across a greater range of MEPs. Indeed, we chose our stimulation intensity as a ‘goldilocks zone’ for this specific research question – i.e., an intensity where participants less responsive to rTMS will not show MEP changes and only people who are very responsive will show expected inhibition. So we delivered cTBS at 80% AMT which, while consistent with the original literature [10], is a lower intensity than other investigators have since employed [50]. We expect that if we stimulated at a higher cTBS intensity, more individuals would respond. Given recent suggestions that many rTMS studies might be under-dosing [17], [51], we strongly urge future studies to test whether stimulating the non-responder group with a higher stimulation intensity can be the solution to eliciting a more uniform response.

We note that our biomarkers are less predictive of the MEP changes at 0 minutes after cTBS. Relatedly, many subjects also show a large change between MEP responses at P0 vs stabilization of effect at later time points P10-P30. This may suggest that there is a need for a waiting period of 10 minutes to see stabilized effects of stimulation. Further, there are baseline differences in average MEP between groups after grouping by IO biomarker – which we corrected for in the linear mixed effects modeling. Yet, the group with higher MEPs at baseline showing a more robust inhibitory response may also be attributable to floor effects among the group with the IO biomarker that also has lower baseline MEPs. Alternatively, this may not be a mechanism but a correlate of the IO biomarker itself in that persons with a higher S_50_ or lower MEP_max_ will also have inherently lower MEPs at baseline because of less responsiveness to stimulation requiring higher stimulation intensities to achieve the same effect.

Our study had some limitations. We only included healthy younger adults in this mechanistic study. Future studies should note that there may be differences across lifespan and across patient populations not captured in this analysis. We suggest that these relationships may be elucidated through use of our MEP_130RMT_ biomarker in future studies. We also acknowledge a small sample size, especially in the context of comparing our IO parameters to the predictions based on BDNF genetic data. While N=21 is a reasonable size for mechanistic rTMS studies, genetic studies generally require much larger samples. We acknowledge that more advanced IO curve fitting models exist, incorporating variability sources beyond what we considered [36], [47]. For example, MEP variability changes across IO curve intensities may more extensively explain the dose-response relationship [52]. However we selected the commonly used Boltzmann sigmoidal IO fitting methodology, as we had largely uniform MEP variances and this classical IO sigmoid curve has well characterized fit parameters in the neuromodulation literature [31], [39]. Additionally, we did not assess IO slopes after cTBS to see if they changed as a result of neuromodulation [17]. Some studies have found that high frequency rTMS applied to the primary motor cortex increases corticospinal excitability causing increased IO slope and reduced motor threshold [30], [53], leading to further questions about the inhibitory and excitatory mechanistic effects on motor cortex cortical networks [44]. We focused on IO curve parameters that have been shown to be stable across days [38] as our study protocol did record IO curves from participants between 1 to 7 days before the cTBS study. Additionally, we only measured MEPs for time points till 30 minutes out from the delivery of cTBS as it is known to affect motor task performance up to this duration in healthy subjects [10]. Future work can extend the length of collected stimulation effects to estimate the duration of predictive capabilities of these IO based parameters.

Our findings may have relevance well beyond the motor cortex, to other rTMS stimulation sites. Future work should investigate whether MEP based IO parameters generalize to other TMS sites and indications across the motor cortex and other stimulation sites. While FDI muscle based IO curves are a popular general marker of excitability and form the basis of how TMS intensity has been decided, we recognize there may be variation across sites [46]. Established methods show that IO curves are possible for sensory pathways with use in disorders of abnormal sensorimotor integration [54], as well as in mapping excitability of any cortical location through TMS-evoked EEG potentials (TEPs) [55] including in real time [56]. Our research then opens a new question for the field as to whether TMS-MEP based IO curves will also predict responses to neuromodulation targeted to other non-motor cortical areas. Given a strong need for stratification in many neuropsychiatric applications of NIBS [1], [2], [3], [18], we suggest future studies should also explore alternative forms of IO curves such as those based upon TMS-EEG TEPs [57], [58] as a more generalizable (albeit more expensive and time consuming) method of customizing to any stimulation site and protocol.

We thus postulate that TMS-MEP IO curves are a strong biomarker of corticospinal excitation, and as we show here, also of responses to cTBS. Based on our research in the context of prior work [29], [46], these IO metrics should also strongly relate to responsiveness to non-invasive neuromodulation in general, including other forms of rTMS, tDCS, and tACS over the motor cortex. We expect that these biomarkers will be directly relevant to stratifying patients to improve outcomes in motor rehabilitation such as after a stroke [59], [60], [61], [62]. We urge the field to adopt prospective reporting on IO curves that will extend and test our current study’s retrospective stratification of patients. The advantages of TBS over other non-invasive brain stimulation strategies are its low intensity, exponentially short duration of application and longer-lasting effects [10], [11]. IO curve based predictors will then allow optimized treatment with the most effective brain stimulation therapies. Neuromodulation protocol specific IO curves may also have expanded uses for predicting responses to other types and locations of rTMS or even NIBS more broadly. The results of our research can help alleviate a common issue around reliability of TBS where a subset of recipients appear to show the opposite neuroplastic effect as intended [42], and may be antidote to recent very discouraging rTMS papers [14], [24], [41]. Our research showing IO curve based biomarkers provide quick, effective stratification into reliable responders or non-responders, opens greater clinical use of TBS, as well as a path to higher efficacy rates of rTMS and numerous other forms of brain stimulation.

## ACKNOWLEDGEMENTS

This work was supported by grants from the NIH and other sources: R01 DC012780-01A1 and the Dana Foundation Brain and Immunoimaging Award to RHH; 5T32NS043126-13 to RW; 5-T32-NS-091006-10 to SYP.

## REFERENCES

[1] K. R. Connolly, A. Helmer, M. A. Cristancho, P. Cristancho, and J. P. O’Reardon, “Effectiveness of Transcranial Magnetic Stimulation in Clinical Practice Post-FDA Approval in the United States: Results Observed With the First 100 Consecutive Cases of Depression at an Academic Medical Center,” J. Clin. Psychiatry, vol. 73, no. 04, pp. e567–e573, Apr. 2012, doi: 10.4088/JCP.11m07413.

[2] R. Voelker, “Brain Stimulation Approved for Obsessive-Compulsive Disorder,” JAMA, vol. 320, no. 11, p. 1098, Sep. 2018, doi: 10.1001/jama.2018.13301.

[3] D. A. Gorelick, A. Zangen, and M. S. George, “Transcranial magnetic stimulation in the treatment of substance addiction,” Ann N Y Acad Sci, vol. 1327, no. 1, pp. 79–93, Oct. 2014, doi: 10.1111/nyas.12479.

[4] T. E. Schlaepfer, M. S. George, and Helen Mayberg on behalf of the WFSBP Task Force on Brain Stimulation, “WFSBP Guidelines on Brain Stimulation Treatments in Psychiatry,” The World Journal of Biological Psychiatry, vol. 11, no. 1, pp. 2–18, Jan. 2010, doi: 10.3109/15622970903170835.

[5] A. T. Sack, “Transcranial magnetic stimulation, causal structure–function mapping and networks of functional relevance,” Current Opinion in Neurobiology, vol. 16, no. 5, pp. 593–599, Oct. 2006, doi: 10.1016/j.conb.2006.06.016.

[6] J. T. Devlin and K. E. Watkins, “Stimulating language: insights from TMS,” Brain, vol. 130, no. 3, pp. 610–622, Mar. 2007, doi: 10.1093/brain/awl331.

[7] J. D. Medaglia et al., “Network Controllability in the Inferior Frontal Gyrus Relates to Controlled Language Variability and Susceptibility to TMS,” J. Neurosci., vol. 38, no. 28, pp. 6399–6410, Jul. 2018, doi: 10.1523/JNEUROSCI.0092-17.2018.

[8] A. Pascual-Leone, J. M. Tormos, J. Keenan, F. Tarazona, C. Cañete, and M. D. Catalá, “Study and Modulation of Human Cortical Excitability With Transcranial Magnetic Stimulation:,” Journal of Clinical Neurophysiology, vol. 15, no. 4, pp. 333–343, Jul. 1998, doi: 10.1097/00004691-199807000-00005.

[9] N. Bolognini, A. Pascual-Leone, and F. Fregni, “Using non-invasive brain stimulation to augment motor training-induced plasticity,” J NeuroEngineering Rehabil, vol. 6, no. 1, p. 8, Dec. 2009, doi: 10.1186/1743-0003-6-8.

[10] Y.-Z. Huang, M. J. Edwards, E. Rounis, K. P. Bhatia, and J. C. Rothwell, “Theta Burst Stimulation of the Human Motor Cortex,” Neuron, vol. 45, no. 2, pp. 201–206, Jan. 2005, doi: 10.1016/j.neuron.2004.12.033.

[11] S. W. Chung, A. T. Hill, N. C. Rogasch, K. E. Hoy, and P. B. Fitzgerald, “Use of theta-burst stimulation in changing excitability of motor cortex: A systematic review and meta-analysis,” Neuroscience & Biobehavioral Reviews, vol. 63, pp. 43–64, Apr. 2016, doi: 10.1016/j.neubiorev.2016.01.008.

[12] B. Hordacre, M. C. Ridding, and M. R. Goldsworthy, “Response variability to non-invasive brain stimulation protocols,” Clinical Neurophysiology, vol. 126, no. 12, pp. 2249–2250, Dec. 2015, doi: 10.1016/j.clinph.2015.04.052.

[13] A.-M. Vallence, M. R. Goldsworthy, N. A. Hodyl, J. G. Semmler, J. B. Pitcher, and M. C. Ridding, “Inter- and intra-subject variability of motor cortex plasticity following continuous theta-burst stimulation,” Neuroscience, vol. 304, pp. 266–278, Sep. 2015, doi: 10.1016/j.neuroscience.2015.07.043.

[14] P. O. Boucher et al., “Sham-derived effects and the minimal reliability of theta burst stimulation,” Sci Rep, vol. 11, no. 1, p. 21170, Oct. 2021, doi: 10.1038/s41598-021-98751-w.

[15] J. F. M. Müller-Dahlhaus, Y. Orekhov, Y. Liu, and U. Ziemann, “Interindividual variability and age-dependency of motor cortical plasticity induced by paired associative stimulation,” Exp Brain Res, vol. 187, no. 3, pp. 467–475, May 2008, doi: 10.1007/s00221-008-1319-7.

[16] M. Inghilleri, A. Conte, A. Currà, V. Frasca, C. Lorenzano, and A. Berardelli, “Ovarian hormones and cortical excitability. An rTMS study in humans,” Clin Neurophysiol, vol. 115, no. 5, pp. 1063–1068, May 2004, doi: 10.1016/j.clinph.2003.12.003.

[17] G. Cotovio et al., “Day-to-day variability in motor threshold during rTMS treatment for depression: Clinical implications,” Brain Stimulation, vol. 14, no. 5, pp. 1118–1125, Sep. 2021, doi: 10.1016/j.brs.2021.07.013.

[18] A. Frick, J. Persson, and R. Bodén, “Habitual caffeine consumption moderates the antidepressant effect of dorsomedial intermittent theta-burst transcranial magnetic stimulation,” J Psychopharmacol, vol. 35, no. 12, pp. 1536–1541, Dec. 2021, doi: 10.1177/02698811211058975.

[19] H. C. Dresang et al., “Genetic and Neurophysiological Biomarkers of Neuroplasticity Inform Post-Stroke Language Recovery,” Neurorehabil Neural Repair, vol. 36, no. 6, pp. 371–380, Jun. 2022, doi: 10.1177/15459683221096391.

[20] S. Parchure et al., “Brain-Derived Neurotrophic Factor Gene Polymorphism Predicts Response to Continuous Theta Burst Stimulation in Chronic Stroke Patients,” Neuromodulation, vol. 25, no. 4, pp. 569–577, Jun. 2022, doi: 10.1111/ner.13495.

[21] P. Shah-Basak et al., “Brain-Derived Neurotrophic Factor Polymorphism Influences Response to Single-Pulse Transcranial Magnetic Stimulation at Rest,” Neuromodulation: Technology at the Neural Interface, vol. 24, no. 5, pp. 854–862, Jul. 2021, doi: 10.1111/ner.13287.

[22] B. Cheeran et al., “A common polymorphism in the brain derived neurotrophic factor gene (*BDNF*) modulates human cortical plasticity and the response to rTMS,” The Journal of Physiology, vol. 586, no. 23, pp. 5717–5725, Dec. 2008, doi: 10.1113/jphysiol.2008.159905.

[23] C. Mastroeni et al., “Brain-Derived Neurotrophic Factor – A Major Player in Stimulation-Induced Homeostatic Metaplasticity of Human Motor Cortex?,” PLoS ONE, vol. 8, no. 2, p. e57957, Feb. 2013, doi: 10.1371/journal.pone.0057957.

[24] J. H. Park, “Reliability of theta burst stimulation as a neuromodulation tool,” Journal of Neurophysiology, vol. 127, no. 6, pp. 1532–1534, Jun. 2022, doi: 10.1152/jn.00507.2021.

[25] Y. Ugawa, “Synaptic and Axonal Plasticity Induction in the Human Cerebral Cortex,” in *Innovative Medicine*, K. Nakao, N. Minato, and S. Uemoto, Eds., Tokyo: Springer Japan, 2015, pp. 295–306. doi: 10.1007/978-4-431-55651-0_24.

[26] D. A. Spampinato, J. Ibanez, L. Rocchi, and J. Rothwell, “Motor potentials evoked by transcranial magnetic stimulation: interpreting a simple measure of a complex system,” The Journal of Physiology, vol. 601, no. 14, pp. 2827–2851, Jul. 2023, doi: 10.1113/JP281885.

[27] E. Ortega-Robles, J. Cantillo-Negrete, R. I. Carino-Escobar, and O. Arias-Carrión, “Methodological approach for assessing motor cortical excitability changes with single-pulse transcranial magnetic stimulation,” MethodsX, vol. 11, p. 102451, Dec. 2023, doi: 10.1016/j.mex.2023.102451.

[28] H. Devanne, B. A. Lavoie, and C. Capaday, “Input-output properties and gain changes in the human corticospinal pathway:,” Exp Brain Res, vol. 114, no. 2, pp. 329–338, Apr. 1997, doi: 10.1007/PL00005641.

[29] T. J. Carroll, S. Riek, and R. G. Carson, “Reliability of the input–output properties of the cortico-spinal pathway obtained from transcranial magnetic and electrical stimulation,” Journal of Neuroscience Methods, vol. 112, no. 2, pp. 193–202, Dec. 2001, doi: 10.1016/S0165-0270(01)00468-X.

[30] E. Houdayer, A. Degardin, F. Cassim, P. Bocquillon, P. Derambure, and H. Devanne, “The effects of low- and high-frequency repetitive TMS on the input/output properties of the human corticospinal pathway,” Exp Brain Res, vol. 187, no. 2, pp. 207–217, May 2008, doi: 10.1007/s00221-008-1294-z.

[31] C. Kemlin et al., “Redundancy Among Parameters Describing the Input-Output Relation of Motor Evoked Potentials in Healthy Subjects and Stroke Patients,” Front. Neurol., vol. 10, p. 535, May 2019, doi: 10.3389/fneur.2019.00535.

[32] S. N. Kukke, R. W. Paine, C.-C. Chao, A. C. De Campos, and M. Hallett, “Efficient and Reliable Characterization of the Corticospinal System Using Transcranial Magnetic Stimulation,” Journal of Clinical Neurophysiology, vol. 31, no. 3, pp. 246–252, Jun. 2014, doi: 10.1097/WNP.0000000000000057.

[33] M. C. Ridding and J. C. Rothwell, “Stimulus/response curves as a method of measuring motor cortical excitability in man,” Electroencephalography and Clinical Neurophysiology/Electromyography and Motor Control, vol. 105, no. 5, pp. 340–344, Oct. 1997, doi: 10.1016/S0924-980X(97)00041-6.

[34] J. C. Keel, M. J. Smith, and E. M. Wassermann, “A safety screening questionnaire for transcranial magnetic stimulation,” Clinical Neurophysiology, vol. 112, no. 4, p. 720, Apr. 2001, doi: 10.1016/S1388-2457(00)00518-6.

[35] C. Capaday, “Neurophysiological methods for studies of the motor system in freely moving human subjects,” Journal of Neuroscience Methods, vol. 74, no. 2, pp. 201–218, Jun. 1997, doi: 10.1016/S0165-0270(97)02250-4.

[36] C. Capaday, “On the variability of motor-evoked potentials: experimental results and mathematical model,” Exp Brain Res, vol. 239, no. 10, pp. 2979–2995, Oct. 2021, doi: 10.1007/s00221-021-06169-7.

[37] F. Hummel and L. G. Cohen, “Improvement of Motor Function with Noninvasive Cortical Stimulation in a Patient with Chronic Stroke,” Neurorehabil Neural Repair, vol. 19, no. 1, pp. 14–19, Mar. 2005, doi: 10.1177/1545968304272698.

[38] J.-M. Therrien-Blanchet et al., “Stability and test–retest reliability of neuronavigated TMS measures of corticospinal and intracortical excitability,” Brain Research, vol. 1794, p. 148057, Nov. 2022, doi: 10.1016/j.brainres.2022.148057.

[39] C. Capaday, B. A. Lavoie, H. Barbeau, C. Schneider, and M. Bonnard, “Studies on the Corticospinal Control of Human Walking. I. Responses to Focal Transcranial Magnetic Stimulation of the Motor Cortex,” Journal of Neurophysiology, vol. 81, no. 1, pp. 129–139, Jan. 1999, doi: 10.1152/jn.1999.81.1.129.

[40] R Core Team, R: A Language and Environment for Statistical Computing. (2024). R Foundation for Statistical Computing, Vienna, Austria. [Online]. Available: https://www.R-project.org/

[41] J. Magnuson et al., “Neuromodulatory effects and reproducibility of the most widely used repetitive transcranial magnetic stimulation protocols,” PLoS ONE, vol. 18, no. 6, p. e0286465, Jun. 2023, doi: 10.1371/journal.pone.0286465.

[42] R. A. Ozdemir et al., “Reproducibility of cortical response modulation induced by intermittent and continuous theta-burst stimulation of the human motor cortex,” Brain Stimul, vol. 14, no. 4, pp. 949–964, 2021, doi: 10.1016/j.brs.2021.05.013.

[43] M. R. Goldsworthy, A.-M. Vallence, N. A. Hodyl, J. G. Semmler, J. B. Pitcher, and M. C. Ridding, “Probing changes in corticospinal excitability following theta burst stimulation of the human primary motor cortex,” Clinical Neurophysiology, vol. 127, no. 1, pp. 740–747, Jan. 2016, doi: 10.1016/j.clinph.2015.06.014.

[44] W. Gerschlager, H. R. Siebner, and J. C. Rothwell, “Decreased corticospinal excitability after subthreshold 1 Hz rTMS over lateral premotor cortex,” Neurology, vol. 57, no. 3, pp. 449–455, Aug. 2001, doi: 10.1212/WNL.57.3.449.

[45] Y.-Z. Huang, R.-S. Chen, J. C. Rothwell, and H.-Y. Wen, “The after-effect of human theta burst stimulation is NMDA receptor dependent,” Clinical Neurophysiology, vol. 118, no. 5, pp. 1028–1032, May 2007, doi: 10.1016/j.clinph.2007.01.021.

[46] L. M. Koponen et al., “Transcranial magnetic stimulation input-output curve slope differences suggest variation in recruitment across muscle representations in primary motor cortex,” Front Hum Neurosci, vol. 18, p. 1310320, 2024, doi: 10.3389/fnhum.2024.1310320.

[47] A. V. Peterchev, S. M. Goetz, G. G. Westin, B. Luber, and S. H. Lisanby, “Pulse width dependence of motor threshold and input–output curve characterized with controllable pulse parameter transcranial magnetic stimulation,” Clinical Neurophysiology, vol. 124, no. 7, pp. 1364–1372, Jul. 2013, doi: 10.1016/j.clinph.2013.01.011.

[48] J. Temesi, M. Gruet, T. Rupp, S. Verges, and G. Y. Millet, “Resting and active motor thresholds versus stimulus-response curves to determine transcranial magnetic stimulation intensity in quadriceps femoris,” J Neuroeng Rehabil, vol. 11, p. 40, Mar. 2014, doi: 10.1186/1743-0003-11-40.

[49] B. Lu, “BDNF and activity-dependent synaptic modulation,” Learn Mem, vol. 10, no. 2, pp. 86–98, 2003, doi: 10.1101/lm.54603.

[50] E. J. Cole et al., “Stanford Accelerated Intelligent Neuromodulation Therapy for Treatment-Resistant Depression,” AJP, vol. 177, no. 8, pp. 716–726, Aug. 2020, doi: 10.1176/appi.ajp.2019.19070720.

[51] D. Cappon, T. Den Boer, C. Jordan, W. Yu, E. Metzger, and A. Pascual-Leone, “Transcranial magnetic stimulation (TMS) for geriatric depression,” Ageing Research Reviews, vol. 74, p. 101531, Feb. 2022, doi: 10.1016/j.arr.2021.101531.

[52] S. M. Goetz, B. Luber, S. H. Lisanby, and A. V. Peterchev, “A Novel Model Incorporating Two Variability Sources for Describing Motor Evoked Potentials,” Brain Stimulation, vol. 7, no. 4, pp. 541–552, Jul. 2014, doi: 10.1016/j.brs.2014.03.002.

[53] E. M. Khedr, J. C. Rothwell, M. A. Ahmed, O. A. Shawky, and M. Farouk, “Modulation of motor cortical excitability following rapid-rate transcranial magnetic stimulation,” Clinical Neurophysiology, vol. 118, no. 1, pp. 140–145, Jan. 2007, doi: 10.1016/j.clinph.2006.09.006.

[54] M. Fischer and M. Orth, “Short-latency sensory afferent inhibition: conditioning stimulus intensity, recording site, and effects of 1 Hz repetitive TMS,” Brain Stimulation, vol. 4, no. 4, pp. 202–209, Oct. 2011, doi: 10.1016/j.brs.2010.10.005.

[55] R. M. Helling et al., “TMS-evoked EEG potentials demonstrate altered cortical excitability in migraine with aura,” Brain Topogr, vol. 36, no. 2, pp. 269–281, Mar. 2023, doi: 10.1007/s10548-023-00943-2.

[56] S. Casarotto et al., “The rt-TEP tool: real-time visualization of TMS-Evoked Potentials to maximize cortical activation and minimize artifacts,” Journal of Neuroscience Methods, vol. 370, p. 109486, Mar. 2022, doi: 10.1016/j.jneumeth.2022.109486.

[57] L. Krile, E. Ensafi, J. Cole, M. Noor, A. B. Protzner, and A. McGirr, “A dose-response characterization of transcranial magnetic stimulation intensity and evoked potential amplitude in the dorsolateral prefrontal cortex,” Sci Rep, vol. 13, no. 1, p. 18650, Oct. 2023, doi: 10.1038/s41598-023-45730-y.

[58] D. Momi, Z. Wang, and J. D. Griffiths, “TMS-evoked responses are driven by recurrent large-scale network dynamics,” eLife, vol. 12, p. e83232, Apr. 2023, doi: 10.7554/eLife.83232.

[59] M. Bowden, “The use of rTMS to augment walking recovery after stroke,” Brain Stimulation, vol. 12, no. 2, pp. 450–451, Mar. 2019, doi: 10.1016/j.brs.2018.12.462.

[60] M. Dafotakis, C. Grefkes, S. B. Eickhoff, H. Karbe, G. R. Fink, and D. A. Nowak, “Effects of rTMS on grip force control following subcortical stroke,” Experimental Neurology, vol. 211, no. 2, pp. 407–412, Jun. 2008, doi: 10.1016/j.expneurol.2008.02.018.

[61] Y. Kim, W. Chang, O. Bang, S. Kim, Y. Park, and P. Lee, “Long-term effects of rTMS on motor recovery in patients after subacute stroke,” J Rehabil Med, vol. 42, no. 8, pp. 758– 764, 2010, doi: 10.2340/16501977-0590.

[62] J.-P. Lefaucheur, “Stroke recovery can be enhanced by using repetitive transcranial magnetic stimulation (rTMS),” Neurophysiologie Clinique/Clinical Neurophysiology, vol. 36, no. 3, pp. 105–115, May 2006, doi: 10.1016/j.neucli.2006.08.011.

