## Supplementary Table 1 for "Predicting Neuroplasticity Effects of Continuous Theta Burst Stimulation with Biomarkers from the Motor Evoked Potential TMS Input-Output Curve"

**SUPPLEMENTAL INFORMATION - for Parchure et. al., 2025**

**Supplemental Table 1:** Individual subjects’ MEP responses to cTBS using 1 way ANOVA for lnMEP ~ Time Point. Values for Time interaction term significance within a subject.

| Subject ID | Sum sq | Mean sq | F-value | p-value | Status |
| --- | --- | --- | --- | --- | --- |
| BME02 | 2.63 | 0.53 | 0.82 | 0.53 | Non-responder |
| BME03 | 0.55 | 0.11 | 0.21 | 0.96 | Non-responder |
| BME04 | 1.65 | 0.41 | 1.63 | 0.17 | Non-responder |
| BME05 | 7.73 | 1.93 | 8.43 | **<0.001** | **Responder** |
| BME06 | 5.64 | 1.41 | 2.47 | **0.04** | **Responder** |
| BME07 | 87.56 | 21.89 | 82.60 | **<0.001** | **Responder** |
| BME08 | 12.79 | 3.20 | 2.70 | **0.03** | **Responder** |
| BME10 | 23.04 | 5.76 | 7.87 | **<0.001** | **Responder** |
| BME11 | 3.17 | 0.79 | 2.55 | **0.04** | **Responder** |
| BME13 | 12.78 | 3.20 | 29.87 | **<0.001** | **Responder** |
| BME14 | 15.11 | 3.78 | 3.74 | **0.006** | **Responder** |
| BME15 | 3.29 | 0.82 | 1.60 | 0.17 | Non-responder |
| BME18 | 6.64 | 1.66 | 4.08 | **0.003** | **Responder** |
| BME20 | 2.87 | 0.72 | 0.69 | 0.60 | Non-responder |
| BME21 | 5.88 | 1.47 | 2.41 | 0.05 | Non-responder |
| BME22 | 218.63 | 54.66 | 146.55 | **<0.001** | **Responder** |
| BME24 | 141.37 | 35.34 | 139.88 | **<0.001** | **Responder** |
| BME25 | 7.04 | 1.76 | 2.11 | 0.08 | Non-responder |
| BME27 | 16.51 | 4.13 | 7.47 | **<0.001** | **Responder** |
| BME28 | 7.55 | 1.89 | 3.11 | **0.017** | **Responder** |
| BME29 | 54.79 | 13.70 | 12.43 | **<0.001** | **Responder** |

Key: Sum sq = sum of squares; Mean sq = mean of squares; Df (degrees of freedom) = 4 for all
